# napariTFM: An Open-Source Tool for Traction Force Microscopy and Monolayer Stress Microscopy

**DOI:** 10.1101/2025.10.14.682385

**Authors:** Artur Ruppel, Dennis Wörthmüller, Martial Balland, François Fagotto

## Abstract

Cellular force generation and transmission are fundamental processes driving cell migration, division, tissue morphogenesis, and disease progression. Traction Force Microscopy (TFM) and Monolayer Stress Microscopy (MSM) have emerged as essential techniques for quantifying these mechanical processes, but current software solutions are fragmented across multiple platforms with varying degrees of usability and accessibility. Here, we present napariTFM, a comprehensive open-source plugin for the napari image viewer that integrates state-of-the-art algorithms for both TFM and MSM analysis within an intuitive graphical user interface. The software implements TV-L1 optical flow for displacement analysis, Fourier Transform Traction Cytometry (FTTC) for force reconstruction, and finite element methods for stress calculation, supporting both single-frame and time-series analysis of 2D microscopy data. Systematic validation using synthetic datasets with known ground truth values demonstrated excellent accuracy, with correlation coefficients above 0.9 for most situations. Real-time parameter adjustment and immediate visualization capabilities enable interactive optimization of analysis parameters and quality assessment during processing. Finally, we demonstrate the software’s capabilities through analysis of optogenetic contractility experiments in cell doublets. napariTFM addresses critical gaps in the cellular mechanics software ecosystem by combining algorithmic rigor with practical usability, providing the research community with an accessible platform for quantitative studies of cellular force generation and transmission.

## 1 Introduction

Cellular force generation and transmission are fundamental processes that drive and regulate critical biological functions including cell migration, division, tissue morphogenesis, and disease progression [1, 2, 3]. Over the past decades, Traction Force Microscopy (TFM) and Monolayer Stress Microscopy (MSM) have emerged as powerful techniques for quantifying these mechanical processes, enabling researchers to measure cell-substrate forces and internal cellular stresses with high spatial and temporal resolution [4, 5, 6].

TFM reconstructs cellular traction forces by measuring substrate deformations caused by adherent cells on elastic substrates embedded with fiducial markers [7]. The technique typically involves comparing images of fluorescent beads in stressed (cell-attached) and relaxed (cell-removed) states, followed by computational reconstruction of force fields. Several computational approaches have been developed for this inverse problem, including Fourier Transform Traction Cytometry (FTTC) [8], Boundary Element Methods (BEM) [9], and finite element approaches [10], each with distinct advantages and limitations depending on the experimental context. MSM extends this analysis by calculating internal stress distributions within cell monolayers [6, 11]. By modeling cellular tissues as thin elastic sheets, MSM enables determination of intercellular force transmission and stress propagation. Important contributions from the biomechanics community have advanced our understanding of these methods’ capabilities and limitations, including developments in 4D TFM [12], quantification of active versus resistive stresses [13], and theoretical frameworks for stress inference in confluent tissues [14].

Despite the widespread adoption of these techniques, significant barriers limit their accessibility to the broader biological research community. Current software solutions are fragmented across multiple platforms with varying degrees of usability, documentation quality, and computational requirements. The TFM software ecosystem includes diverse implementations: ImageJ plugins such as Qingzong Tseng’s PIV and FTTC implementations [15], JEasyTFM [16] and iTACS [17], the stand-alone tool Cellogram [18] for reference-free real-time analysis, MATLAB tools such as TFMLAB [19] for 4D TFM capabilities or *μ*-inferforce [9], implementing both FTTC and BEM algorithms, and Python tools such as pyTFM [20] and our own previous tool batchTFM[21]. For MSM analysis, pyTFM and iTACS are, to our knowledge, the only freely available solutions.

However, several challenges remain for researchers seeking to implement these techniques. First, the fragmented software landscape requires users to switch between different platforms and programming environments for complete analysis workflows. Second, selecting appropriate algorithmic parameters (regularization values, mesh densities, filtering parameters) requires significant expertise and often lacks real-time visual feedback during optimization. Third, processing time-series datasets at scale demands robust batch processing capabilities that many existing tools lack. Finally, researchers without programming expertise face substantial barriers to entry, limiting the techniques’ adoption despite their biological value.

To this end, we developed napariTFM, a comprehensive TFM/MSM napari plugin that addresses these critical gaps through an intuitive graphical user interface, immediate feedback on parameter selection effects, and powerful batch processing capabilities for high-throughput experiments. The plugin leverages the Python-based napari image viewer [22], which is emerging as a powerful platform for biological image analysis with active community development and an extensive plugin ecosystem for comprehensive microscopy workflows.

The plugin supports both single-frame and time-series analysis of 2D microscopy images. napariTFM implements state-of-the-art algorithms including TV-11 optical flow for displacement analysis, FTTC for force reconstruction, and finite element methods for stress calculation.

## 2 Methods

### 2.1 Assumptions and Limitations

napariTFM addresses the methodological complexities of force microscopy by making established computational approaches accessible, transparent, and easy to use for biologists conducting diverse experiments. The plugin implements well-established algorithms including TV-11 optical flow for displacement analysis [23], FTTC for force reconstruction [8, 10], and finite element methods for stress calculation [6, 20]. We chose these specific methods based on their favorable balance of computational efficiency, robustness to noise, and ability to handle the range of displacement magnitudes encountered in typical biological experiments. However, we acknowledge that alternative approaches may be preferable in specific contexts. These methods rely on fundamental assumptions about substrate and cellular material properties (linear elasticity, homogeneity), measurement conditions (2D imaging of inherently 3D systems), and force balance (which may not hold locally even when satisfied globally). Different algorithmic approaches handle these challenges differently, with trade-offs between computational efficiency, noise robustness, and accuracy under various experimental conditions [7, 10]. Important considerations for users include the effects of spherical aberration at image edges in large monolayers, and the current implementation of boundary conditions for MSM which work only when cell borders are clearly visible. Future versions will address boundary conditions for confluent layers without visible borders and could incorporate multiple algorithmic options with guidance for users on method selection.

### 2.2 Software Architecture and lmplementation

napariTFM is implemented as a Python 3.6+ package designed to integrate with the napari image viewer as an optional plugin. The complete analysis workflow from raw input data through preprocessing, displacement analysis, force calculation, and stress analysis is outlined in Figure 1. The software provides both a graphical user interface through napari (Figure 2) and a standalone Python library for programmatic access. The core computational components utilize established opensource packages including NumPy for array operations, SciPy for scientific computing, OpenCV for image processing, scikit-image for automated detection and drift correction, and matplotlib for visualization. The plugin organizes data using a hierarchical structure where each experimental field of view is represented as a frame containing multiple image layers. Input images (substrate in tensed and relaxed states, and optional cell images) are stored as separate layers, with analysis outputs added as additional layers during processing. Cell masks are created through simple threshold segmentation of cell images or can be provided externally as TIFF files.

**Figure 1:**
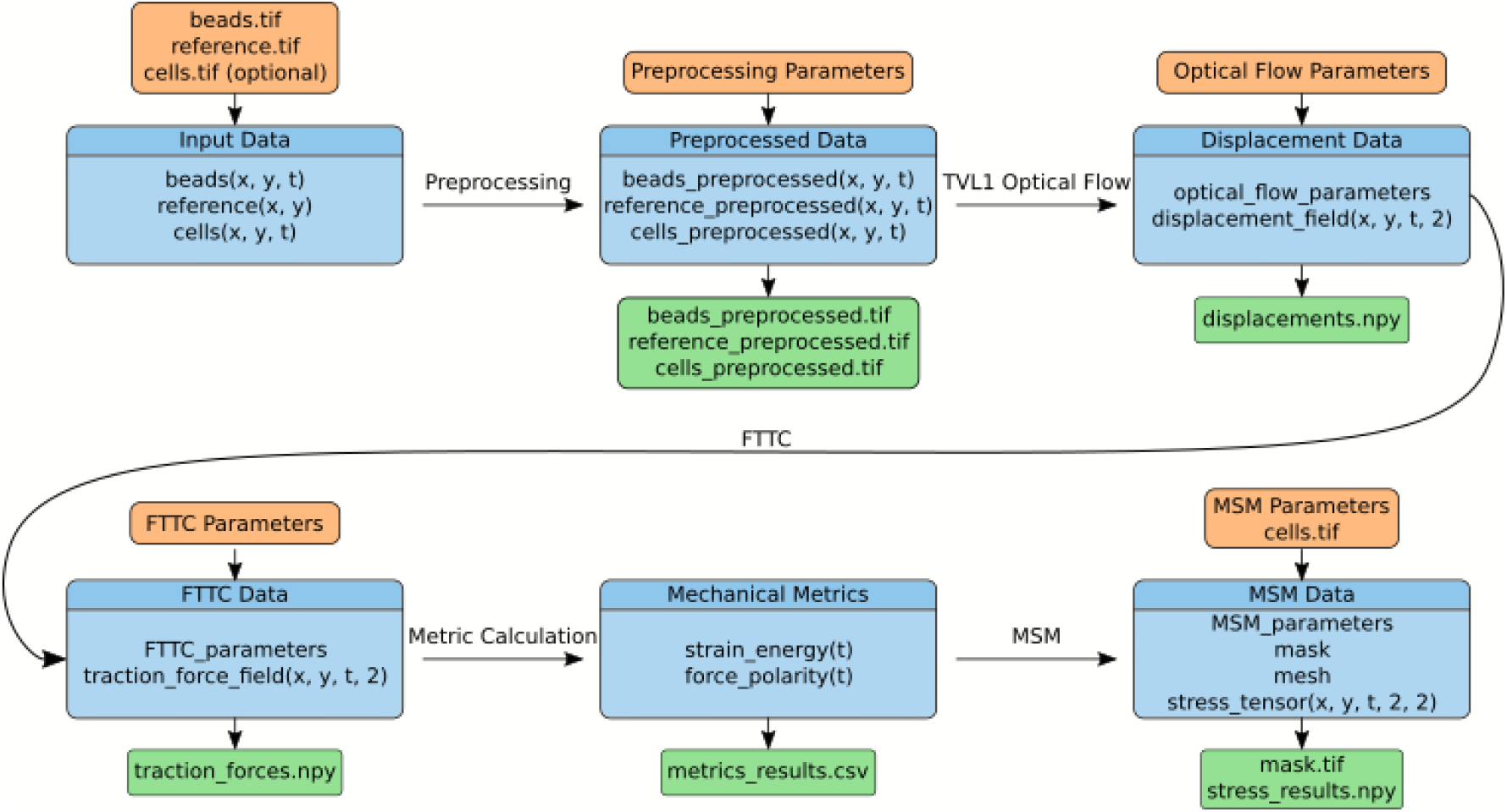
napariTFM workflow and data structure. Schematic overview of the napariTFM analysis pipeline showing the flow from raw input data (beads, reference, and optional cell images) through preprocessing, displacement analysis using TV-L1 optical flow, force calculation via FTTC (Fourier Transform Traction Cytometry), and stress analysis using MSM (Monolayer Stress Microscopy). Orange boxes indicate input data, blue boxes show analysis steps and their internal data structures, and green boxes indicate output files generated at each step.

**Figure 2:**
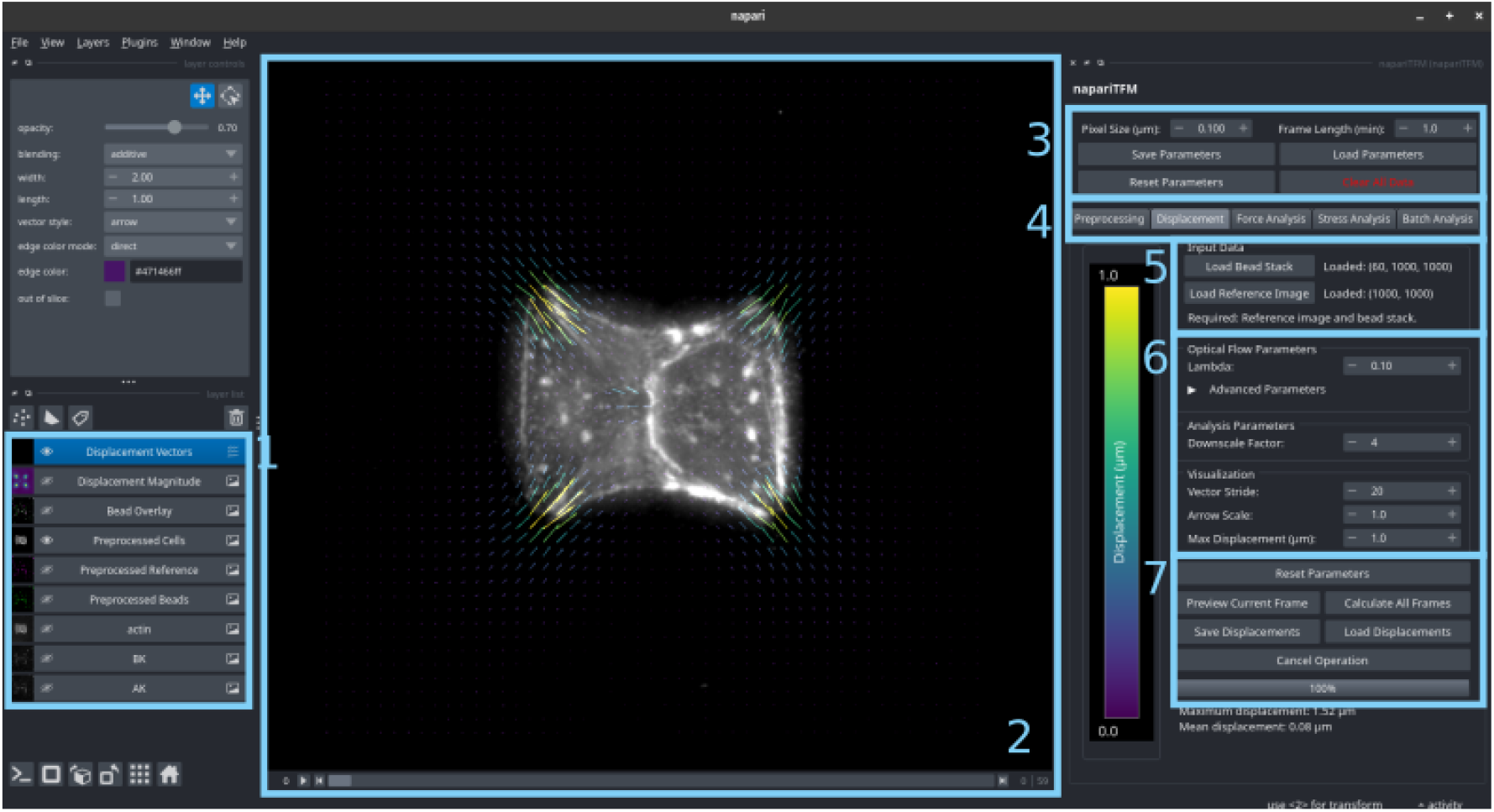
napariTFM user interface in napari. Screenshot of the napariTFM plugin integrated within the napari image viewer, showing: (1) the main image display with fluorescent bead data and overlaid displacement vectors, (2) napari s layer controls, (3) global parameters panel, (4) tabs that access controls for different analysis steps, (5) data input panel, (6) local parameters for the selected step, and (7) action controls (analyze, save data, preview etc.). The interface provides real-time parameter adjustment and visualization of results.

### 2.3 Displacement Field Calculation

Substrate deformations are calculated using the TV-11 optical flow algorithm [24, 25], which is particularly well-suited for TFM analysis due to its ability to handle steep gradients in displacement fields, large displacements through multi-scale analysis, and provision of subpixel accuracy while being robust to intensity variations. The inherent smoothness constraint in TV-11 optical flow makes it particularly robust to imperfect experimental conditions such as bead aggregates or non-uniform bead densities, reducing the need for post-processing steps like outlier removal that are sometimes necessary with other displacement tracking methods. However, users should be aware that this smoothness constraint can also over-regularize displacement fields in regions with genuinely high deformation gradients.

The TV-11 algorithm minimizes an energy functional that combines brightness constancy assumptions (beads maintain intensity) with total variation regularization (smooth displacement fields) and additional constraints for numerical stability. It uses a multi-scale pyramid approach where images are analyzed at different resolution levels. Large displacements are captured at coarse scales while fine details are refined at higher resolutions. Key algorithm parameters include lambda (*λ*) which controls the balance between data fitting and smoothness (typical values 0.01-1.0, with lower values providing more smoothing for noisy data and higher values preserving detail for clear images), pyramid scales (number of resolution levels, typically 3-5), warps (number of iterative refinements per scale), epsilon (stopping criterion for optimization), and scale step (factor between pyramid levels, typically 0.5-0.8). Global image drift correction is performed using phase cross-correlation of the entire image pair, followed by image alignment and cropping to the overlapping field of view.

### 2.4 Traction Force Reconstruction

Traction forces are computed using the Fourier Transform Traction Cytometry (FTTC) method, using the implemtation by Blumberg et al. [10]. FTTC is computationally efficient due to the Fast Fourier Transform, typically processing images in seconds, making it well-suited for batch processing of large time-series datasets.

Linear elasticity theory relates substrate deformations (**u**) and cellular tractions (**t**) through:

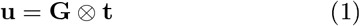

where **G** represents the Green’s tensor for a linearly elastic substrate and ⊗ denotes convolution. By exploiting the convolution theorem, this relationship simplifies in Fourier space to:

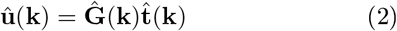

where hats indicate Fourier transforms. The Green’s tensor for a semi-infinite elastic substrate follows the Boussinesq equations, with optional corrections for finite substrate thickness when analyzing larger cell patches.

For regularization, napariTFM uses automatic Generalized Cross-Validation (GCV) to find the optimal regularization parameter *λ* when auto-GCV is selected. When manual regularization is chosen, the user can specify the regularization parameter directly. The FTTC algorithm uses Tikhonov regularization to solve the ill-posed inverse problem of reconstructing forces from displacement measurements. When comparing datasets quantitatively, it is essential to maintain consistent regularization parameters to ensure that differences in recovered forces reflect biological variation rather than analysis artifacts. However, if substrate rigidity differs between conditions, the regularization parameter should be adjusted accordingly as the optimal regularization depends on the mechanical properties of the system. A Lanczos filter with user-defined exponent is applied for additional noise reduction, with higher exponents providing stronger smoothing at the cost of spatial detail. The choice of Lanczos exponent, like the regularization parameter, should remain consistent within comparative studies.

### 2.5 Monolayer Stress Microscopy Implementation

Stress fields in cell monolayers are calculated using the Monolayer Stress Microscopy algorithm developed by Tambe et al. [6], using the implementation from the pyTFM package [20].

The cell monolayer is modeled as a two-dimensional linear elastic sheet in contact with the matrix, where external tractions are balanced by internal stresses.

In the absence of inertial forces, tractions and stresses are balanced according to:

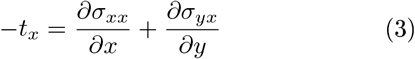

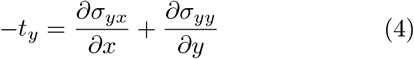

where *σ*_*xx*_, *σ*_*yy*_ are normal stresses in *x*- and *y*-directions, *σ*_*yx*_ is the shear stress, and *t*_*x*_, *t*_*y*_ are traction components.

This differential equation is solved using a finite element method where the cell patch is modeled using triangular mesh elements generated with gmsh. Gmsh provides user-controllable mesh parameters including density and mesh algorithms and also provides mesh quality metrics. Each node is loaded with forces equal in magnitude but opposite in direction to the local tractions. Nodal displacements are calculated by solving:

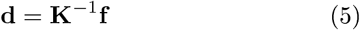

where **d** are nodal displacements, **f** are nodal forces, and **K**^−1^ is the inverse stiffness matrix. The nodal displacements are converted to strains by taking spatial derivatives, and stresses are calculated using the stress-strain relationship for a linearly elastic two-dimensional material. FEM calculations are performed using the SolidsPy Python package.

The FEM algorithm assumes zero net forces and torques on the cell patch. Since TFM ensures global but not local force balance, unbalanced forces and torques are corrected:net forces are removed by subtracting the sum of all force vectors from each node, and net torques are corrected by rotating all force vectors by small angles until zero torque is achieved.

Zero rigid translation and rotation constraints are applied to make the system uniquely solvable. Rather than fixing individual nodes, constraints are formulated globally as:

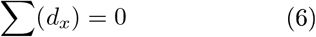

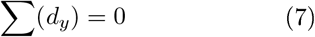

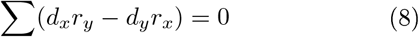

where *r*_*x*_, *r*_*y*_ are distance components from nodes to the grid center of mass. These constraints are incorporated into the system and solved numerically using least-squares minimization.

The Young’s modulus is set to 1 Pa and Poisson’s ratio to 0.5, as the stress calculation is independent of the actual Young’s modulus value and only negligibly influenced by Poisson’s ratio [11]. The triangular mesh density and quality can be adjusted through user-defined parameters, allowing optimization for different cell geometries and experimental and computational requirements.

### 2.6 Synthetic Data Generation for Validation

To validate the accuracy of napariTFM’s displacement analysis, traction force reconstruction, and stress calculation algorithms, we generated synthetic datasets with known ground truth values for systematic comparison.

For traction force microscopy validation, we utilized the DirectMethod repository from Blumberg et al. [10], which provides a forward simulation approach to generate substrate displacements from known force fields. We programmed two force dipoles with identical centers and 90-degree rotation relative to each other, mimicking the characteristic force field pattern of a cell doublet on H-shaped micropatterns, similar to the experimental conditions described in our previous work [26]. The synthetic force fields were designed with varying magnitudes (low, medium, and high) to test algorithm performance across different force scales. To create realistic synthetic bead images, we used OpenCV to deform experimental fluorescent bead images according to the displacement maps generated by the forward simulation.

For monolayer stress microscopy validation, we employed two complementary approaches. First, we used an analytically solved problem consisting of a square plate under uniform loading, where both the internal stress distribution (constant within the plate) and boundary forces are known exactly, providing a rigorous benchmark for algorithm accuracy.

Second, we employed a finite element method (FEM), inspired by the modeling approach in [27, 28] to generate realistic stress maps and corresponding traction force distributions that capture the mechanical behavior of migrating cells. This approach enables validation under more complex and biologically relevant conditions. In this framework, cells are modeled as a two-dimensional active elastic solid via

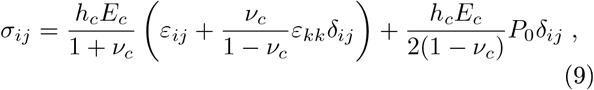

where the first term represents the constitutive relation of a linear elastic material, with *σ*_*ij*_ and *ε*_*ij*_ denoting the Cauchy stress and strain tensor, respectively. The second term introduces an active contractile stress with constant contractility *P*_0_. The cell layer is further characterized by the Young’s modulus *E*_*c*_, Poisson’s ratio *ν*_*c*_, and effective contractile thickness *h*_*c*_. To increase geometric and mechanical complexity, the cell is assumed to adhere at 19 distinct adhesive islands positioned near the periphery similar to [29]. Force balance is then given by

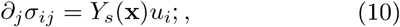

where *Y*_*s*_(**x**) ≠ 0 at adhesion sites (and vanishes otherwise), describing the spring stiffness of the elastic substrate, and **u** is the substrate displacement field [30]. For the parametrization of this continuum model, we closely follow the parameters listed in the Supporting Information of [29]. In our FEM simulations, an initially round cell undergoes isotropic contraction until the force balance in Equation 10 is reached. The resulting cell shape and internal stress patterns are shown in Figure 4C (upper left row).

## 3 Results

We systematically validated napariTFM’s performance across all maJor analysis components using synthetic datasets with known ground truth values, followed by demonstration of its capabilities on real experimental data. The validation encompassed displacement field calculation, traction force reconstruction, and monolayer stress microscopy analysis, confirming the software’s accuracy and reliability for quantitative cellular force measurements.

### 3.1 Displacement Analysis Performance

The displacement field reconstruction achieved excellent accuracy across all tested scenarios (Figure 3A), with correlation coefficients between calculated and ground truth displacement of ≈0.95 for low displacement scenarios and above ≈0.98 for medium and high displacement conditions, demonstrating robust performance across biologically relevant deformation magnitudes.

**Figure 3:**
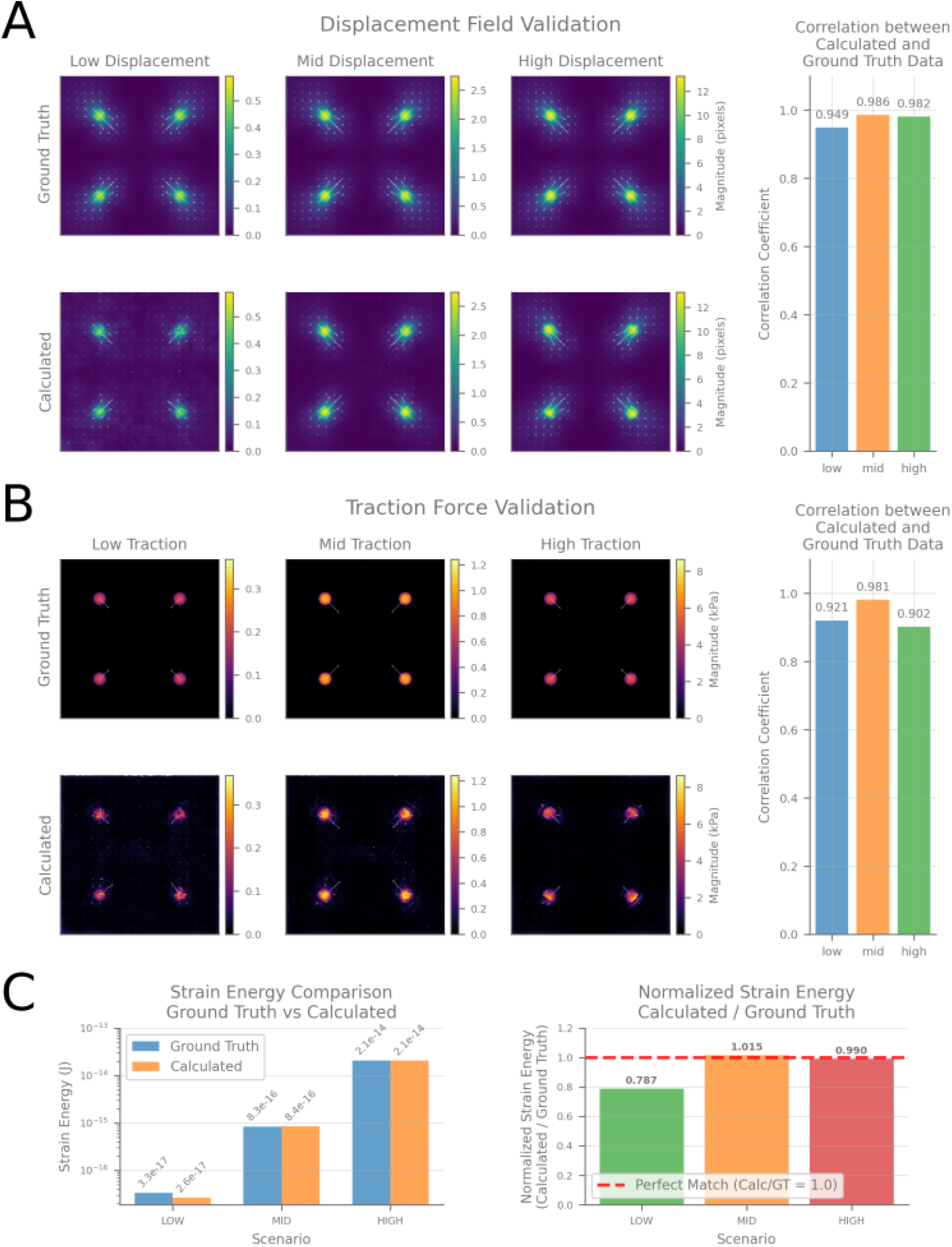
Validation of displacement analysis and traction force calculation. (A) Displacement field validation across three scenarios (low, mid, high displacement magnitudes). Top row shows ground truth displacement fields, bottom row shows calculated results using napariTFM’s TV-11 optical flow algorithm. Vector overlays indicate displacement direction and magnitude. Bar chart shows correlation coefficients between calculated and ground truth data. (B) Traction force validation using FTTC algorithm. 1ayout as in (A), with ground truth (top) and calculated (bottom) traction fields. (C) Strain energy analysis comparing ground truth versus calculated values (left) and normalized ratios (right). The dashed red line indicates perfect agreement (calculated/ground truth = 1.0).

### 3.2 Traction Force Reconstruction Accuracy

Force reconstruction showed high fidelity to ground truth data (Figure 3B), with correlation coefficients ranging from 0.9 to 0.98 across different force magnitudes. Strain energy analysis (Figure 3C) revealed excellent quantitative agreement for medium and high displacement scenarios, with normalized strain energy ratios clustering tightly around the ideal value of 1.0. The low displacement scenario showed reduced accuracy (≈0.79), representing a limiting case where signal-to-noise ratio challenges affect force magnitude recovery.

### 3.3 Monolayer Stress Microscopy Performance

MSM validation using analytical solutions (Figure 4A,B) achieved correlation coefficients above 0.98 for all stress tensor components (*σ*_*xx*_, *σ*_*yy*_, *σ*_*normal*_), with normalized stress ratios very close to 1. FEM simulation validation (Figure 4C,D) with realistic cell-like geometries yielded correlation coefficients between 0.85-0.91, confirming robust performance under complex stress distributions.

**Figure 4:**
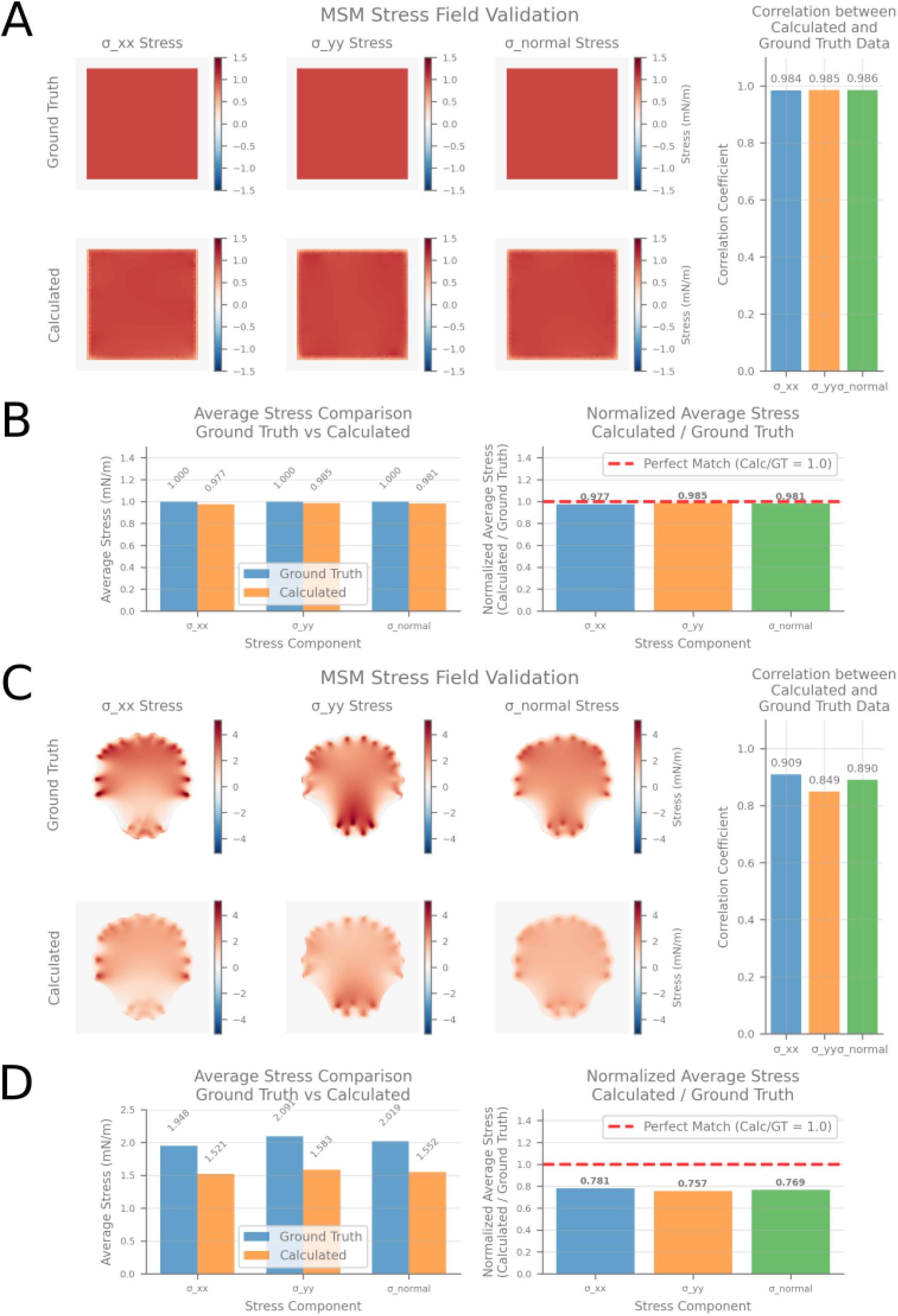
Validation of Monolayer Stress Microscopy (MSM) analysis. (A) Square plate analytical validation using a simple geometric test case with known analytical solution. Stress tensor components (*σ*_*xx*_, *σ*_*yy*_, *σ*_*normal*_) are shown for ground truth (top) and calculated (bottom) fields. Bar chart shows correlation coefficients between calculated and ground truth stress components. (B) Average stress comparison and normalized ratios for square plate validation. The dashed red line indicates perfect agreement (calculated/ground truth = 1.0). (C) FEM-simulation validation using computational data from a finite element model that resembles a snapshot of a migrating cell. Layout as in (A). (D) Average stress comparison and normalized ratios for FEM-simulation validation.

### 3.4 Example Application:Optogenetic Stimulation of Cell Contractility in Cell Doublets

Time-series analysis of optogenetic contractility experiments demonstrated napariTFM’s capability for dynamic biological applications (Figure 5). The experimental data was previously published [26] and reanalyzed here using napariTFM to demonstrate the software’s capabilities. Following stimulation, traction force magnitude increased dramatically for the duration of stimulation and relaxed again after approximately 20 minutes. MSM analysis revealed corresponding changes in internal stress distributions, showing how contractile forces propagate spatially through the cell doublet.

**Figure 5:**
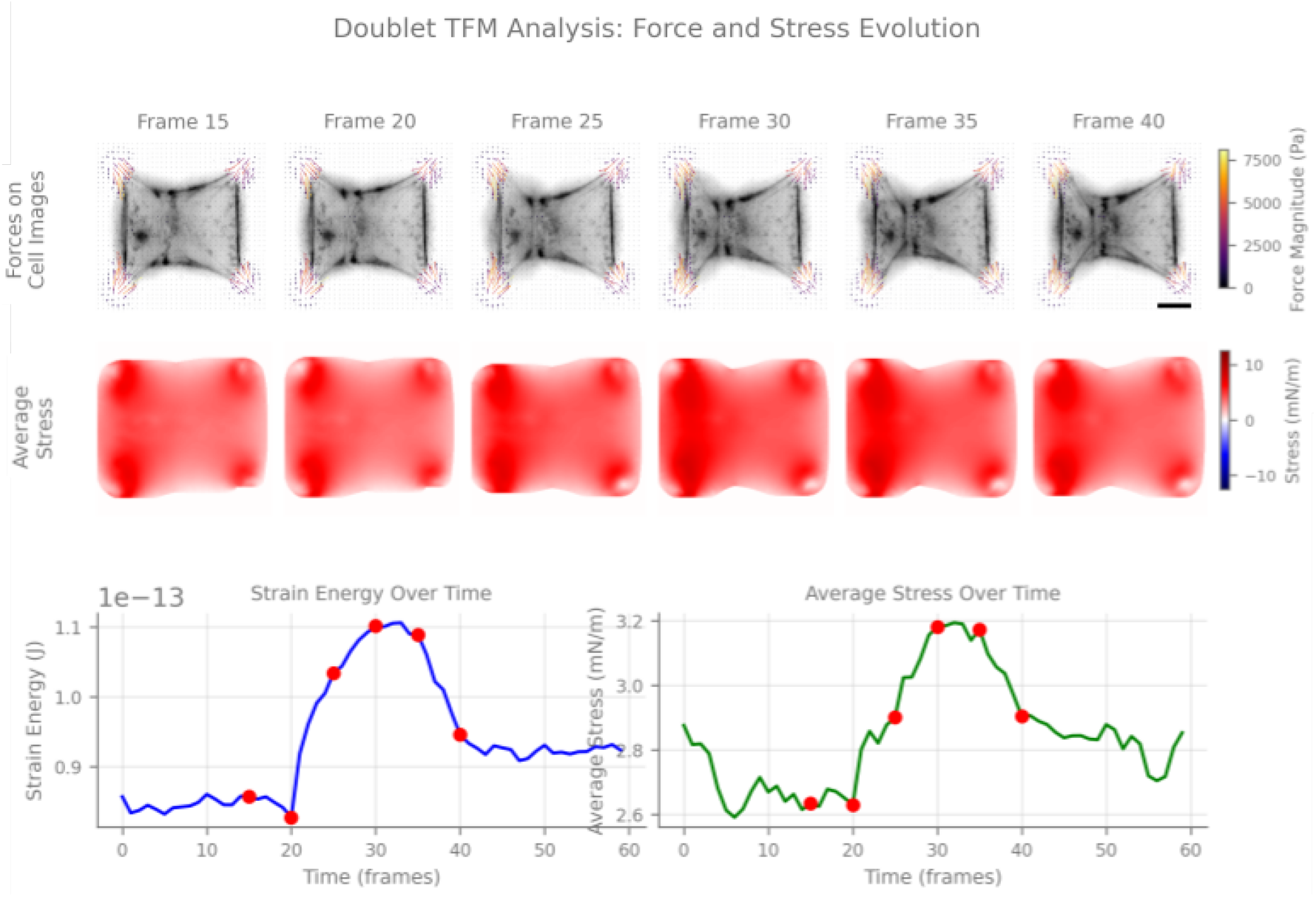
Cell doublet TFM analysis demonstrating temporal evolution of forces and stress. Raw microscopy data from [26] was reanalyzed using napariTFM. Top row shows traction force vectors overlaid on cell images at six time points, with force magnitude color-coded. Middle row displays corresponding average normal stress fields calculated using MSM, showing the spatial distribution of internal cellular stresses. Bottom row presents quantitative time-series analysis:strain energy evolution (left) and spatially-averaged stress over time (right), with red dots highlighting the selected visualization time points. Scale bar:10 µm.

## Discussion

napariTFM addresses a critical need in the cellular mechanics community by providing a comprehensive, userfriendly platform that integrates state-of-the-art algorithms for both traction force microscopy and monolayer stress microscopy. Our validation results demonstrate that the software achieves excellent accuracy across the full range of biologically relevant force and displacement magnitudes, with correlation coefficients consistently above 0.9 for most scenarios.

The integration within the napari ecosystem represents a significant advantage for the broader microscopy community, leveraging napari’s established plugin architecture and intuitive interface design. The real-time parameter adjustment and immediate visualization capabilities address a common limitation of batch-processing tools, allowing users to optimize analysis parameters interactively and assess data quality during processing.

Our synthetic data validation revealed important performance characteristics across different experimental conditions. The reduction in accuracy for low displacement scenarios (normalized strain energy ratio of 0.79) reflects the fundamental signal-to-noise limitations inherent to TFM analysis. This finding is consistent with previous studies and highlights the importance of experimental design considerations, such as substrate stiffness selection and imaging quality optimization, for achieving optimal force measurement accuracy.

While napariTFM’s primary contribution lies in integrating established methods into an accessible platform, it is important to acknowledge the algorithmic choices and their limitations. Our validation demonstrates the accuracy of specific implementations:TV-11 optical flow, FTTC, and finite element MSM. We selected these methods for their favorable balance of computational efficiency, noise robustness, and ability to handle typical biological displacement magnitudes, but alternative approaches (e.g., PIV, BEM) may offer advantages in specific contexts [9, 7]. Future versions could incorporate multiple algorithmic options with guidance for method selection based on experimental requirements.

All force microscopy methods rely on assumptions that users must understand when interpreting results. TFM assumes linear elastic substrate behavior of the substrate to relate measured displacements to cellular traction forces, which means that substrates with more complex material properties such as matrigel or collagen gels, require different methods. For MSM, while the formulation invokes elastic sheet theory, the calculated stress distributions are largely independent of the assumed elastic modulus and only negligibly influenced by Poisson’s ratio [11], making the elasticity assumption less restrictive than it may appear. Standard 2D analysis captures projections of 3D mechanical systems1 potentially missing out-of-plane forces. Regularization in FTTC represents critical trade-offs between noise suppression and spatial detail preservation; napariTFM1s real-time visualization allows interactive assessment1 but optimal parameters ultimately require biological judgment about expected force patterns. MSM requires force balance corrections because TFM ensures global but not local equilibrium. The magnitude of these corrections can serve as a quality metric for the analysis.

Important practical considerations include distinguishing between isolated cells and confluent monolayers1 which require different analytical approaches and boundary conditions. Our current MSM implementation works only when cell borders are clearly defined. Boundary conditions for truly confluent tissues without visible cell-cell boundaries will be addressed in future versions. Additionally1 spherical aberration at image edges and photobleaching in time-lapse experiments can affect displacement accuracy1 emphasizing that careful experimental design remains essential for high-quality force microscopy regardless of computational methods.

Current limitations include the restriction to 2D analysis and the computational requirements for large-scale time-series datasets. Future developments will focus on extending the framework to 2.5D and 3D traction force microscopy1 implementing GPU acceleration for improved processing speed1 and developing specialized tools for high-throughput screening applications.

napariTFM fills an important gap in the TFM and MSM software ecosystem by combining an interactive and user-friendly graphical interface with state-of-theart algorithms. Its open-source nature and integration within the napari platform position it as a valuable resource for advancing quantitative studies of cellular force generation and transmission across diverse biological systems.

## 5 Code Availability

napariTFM is open-source software available at https://github.com/ArturRuppel/napariTFM (GNU General Public License) and can be installed by following the installation procedure outlined in the readme. Synthetic validation datasets and analysis scripts are included in the repository.

## Acknowledgments

We acknowledge the foundational algorithmic contributions that enabled this work:Andreas Bauer and Ben Fabry for the pyTFM implementation of monolayer stress microscopy and finite substrate thickness corrections, and Johannes Blumberg and Ulrich Schwarz for the FTTC implementation and synthetic data generation methods. napariTFM builds upon these established frameworks to provide an integrated analysis platform for the cellular mechanics community. We further would like to thank François Graner for providing valuable feedback on the manuscript.

During the development of this work, we used Claude (Anthropic) as a coding assistant for software development and to improve the clarity and organization of the manuscript text. We reviewed and edited all AI-generated content and code, and take full responsibility for the software implementation and final manuscript.

This work was supported by the ANR (Agence Nationale de la Recherche) grant Inter-s-cal (ANR-21CE13-0042), coordinated by Francois Fagotto. Dennis Worthmiiller received funding from a European Research Council (ERC) grant ERC-SyG 101071793.

